# Intergenerational control of ribosomes under dietary restriction

**DOI:** 10.1101/2025.03.01.640961

**Authors:** Sigma Pradhan, Klement Stojanovski, Joel Tuomaala, Nicholas Stroustrup, Benjamin D. Towbin

## Abstract

Cells adjust their proteome to their environment. Most prominently, ribosome expression scales near linearly with the cellular growth rate to provide sufficient translational capacity while preventing metabolically wasteful ribosomal excess. In microbes, such proteome adjustments can passively perpetuate through symmetric cell division. However, in animals, a passive propagation is hindered by the separation between soma and germline. This separation raises the crucial question whether the proteome of animals is reset at every generation or can be propagated from parent to offspring despite this barrier. We addressed this question by exploring the intergenerational effects of dietary restriction in *C. elegans*, combining proteomics and live imaging. While most proteins showed no intergenerational regulation, ribosomal proteins remained reduced in offspring after maternal dietary restriction. When offspring of dietarily restricted mothers were raised under improved dietary conditions, this reduced ribosome content delayed their growth until normal ribosomal protein levels were restored. Soma-specific maternal inhibition of mTORC1 signalling replicated these effects, while other growth-reducing perturbations, such as reduced insulin signalling or maternal ribosome depletion, did not impact offspring ribosomes. Thus, mTORC1 signalling bridges across the soma-germline divide to regulate ribosome levels of the next generation, likely priming the offspring for the anticipated demand in protein synthesis.

## Introduction

Cells vary the allocation of their proteome resources to different tasks depending on their growth rate^1,2^. When growth is rapid, cells must highly express the protein synthesis machinery, including ribosomes, whereas maintaining such high ribosome expression in growth-limiting environments would be metabolically inefficient.

In bacteria, the cellular ribosome content follows a near linear relation with the growth rate – a phenomenon termed the “growth law” – which is thought to optimize growth across many conditions^3,4^. This relationship between growth and ribosome expression extends at least qualitatively to multicellular animals. At a cellular scale, ribosome expression varies by 3- to 10-fold among human tissues based on their anabolic demands^5,6^. Similarly, at the organismal scale, ribosome expression often scales with the growth rate supported by the environment. For example, under condition of dietary restriction, where growth is slow, the nematode *C. elegans* reduces its ribosome expression accordingly^7^.

Although the regulation of ribosomes in accordance with the growth rate is conserved between multicellular and unicellular forms of life, there is a fundamental difference in the propagation of ribosomes, and the proteome in general, across generations. In microbes, the proteome - and any adjustments to its content – can, in principle, be stably maintained across generations by the equal distribution of proteins between daughter cells during division. In multicellular animals, however, the separation of soma and germline, called the “Weismann barrier”^8^, restricts the passive transmission of the proteome to subsequent generations.

Despite this barrier, the parental environment can have a pronounced impact on the physiology of the offspring^9^, called intergenerational inheritance when lasting for only one generation, or transgenerational inheritance when lasting for longer^10^. Examples include maternal effects on resistance to pathogens^11^, osmotic stress^12^, temperature^13^, or starvation^14^.

While many studies have searched for regulatory signals that sustain an altered gene expression profile in response to maternal environments, such as small RNAs or histone and DNA modifications^10,15–17^, a simpler yet crucial question has remained poorly explored: To what extent is the global partitioning of the proteome of newborn animals, and specifically its allocation to protein translation and ribosomes, determined by the environment and growth rate of their parents? Do animals, like microbes, quantitatively inherit their mother’s proteome allocation to essential cellular machinery in accordance to their growth rate, or is the proteome globally reset between generations and independent of the parental environment?

We investigated these questions by examining the intergenerational effects of dietary restriction (DR) in *C. elegans*, a model system in which the proteomic and physiological response to diet within a generation is well established^7,18,19^. We show that while DR profoundly alters the proteome within a generation, the proteome of the next generation is much less impacted by maternal diet. However, ribosomal proteins represent a notable exception to this rule and maintain reduced levels also in offspring of dietarily restricted mothers. This reduced expression of ribosomes has substantial physiological consequences: when progeny of DR animals was exposed to *ad libitum* (AL) feeding, they displayed a delay in growth and development until they had synthesised normal ribosome levels. Maternal soma-specific inhibition of mTOR signalling – but not of other growth-inhibiting interventions – replicated the effect of DR, suggesting that the ribosome content at hatch is regulated by metabolic signalling from the soma to the germline. Together, our study reveals an important role of intergenerational ribosome control in determining growth performance across generations.

## Results

### Dietary restriction globally alters the proteome within a generation

To determine how dietary conditions alter the protein composition of *C. elegans* within and across generations, we employed tandem-mass-tag (TMT) proteomics. Animals were grown in liquid culture using *E. coli* HB101 as a food source. Dietary restriction (DR) was imposed by a five-fold food dilution compared to *ad libitum* (AL) feeding (2*10^8^ vs. 10^9^ cfu/ml), which slowed down development from three to five days per generation (Fig. 1A).

**Figure 1.**
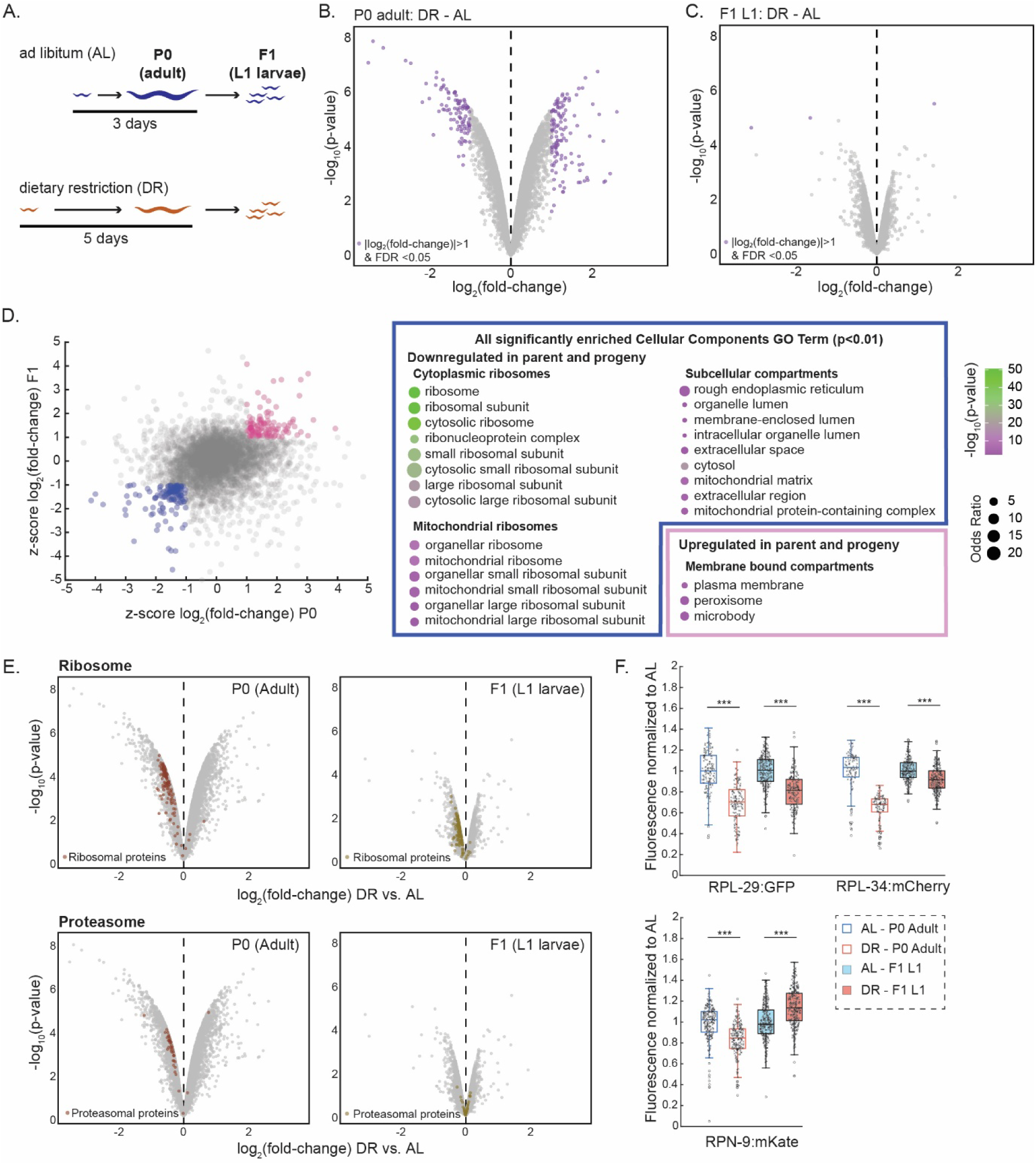
Maternal dietary restriction (DR) reduces ribosome expression in progeny. **(A)** Experimental design of maternal DR. Animals were grown from L1 larvae to adulthood under *ad libitum* (AL) feeding (3 days) or DR (5 days). Adults and their F1 progeny were collected as synchronized L1 larvae for TMT proteomics in n = 3 biological repeats. **(B)** Volcano plot of relative changes in protein levels between DR and AL adults. Purple dots indicate proteins with a |log_2_(fold change)| > 1 at a false discovery rate (FDR) < 0.05. **(C)** As (B), but for L1 larvae of F1 progeny. **(D)** Left: Scatter plot shows consistently upregulated (pink) or downregulated (blue) proteins after DR in parents and their progeny (|z-score| > 1). Right: All cellular component Gene Ontology terms enriched among consistently up or down regulated proteins with odds ratio > 1.5 and adjusted p-value < 0.01 (Benjamini-Hochberg correction). Circle size: odds ratio, Circle colour: significance **(E)** As (B) and (C) but with ribosomal and proteasomal proteins highlighted in colour. **(F)** Quantification of fluorescence intensity (fluorescence per pixel) of indicated endogenously tagged proteins of parents exposed to DR or AL and their L1 progeny by live imaging. RPL-29 and RPL-34 are ribosomal proteins, RPN-9 is a proteasomal protein. Data are normalized to the respective AL conditions. central line: median, box: interquartile range (IQR), whisker: ranges except extreme outliers (>1.5*IQR), individual values: crosses, extreme outliers: circles. For each condition a total of at least n > 110 individuals (for P0) and at least n > 230 individuals (for L1 progeny) were measured over at least m = 3 days. See Supplemental Table S2 for precise sample size and p-values. *** indicate p<10^−10^ (Wilcoxon rank-sum test).

For both dietary conditions, we collected animals at young adulthood stage (containing 1 to 3 eggs), and as synchronised, arrested L1 larvae of the next generation in triplicate repeats. By TMT proteomics, we detected over 6’400 proteins across all replicates and conditions (Fig. S1A). DR substantially altered protein expression in young adults with 104 proteins significantly downregulated, and 125 proteins significantly upregulated by more than 2-fold (Fig. 1B, FDR < 0.05, Supplemental Table S1a). Downregulated proteins were enriched for translational machinery, particularly in ribosomes and vitellogenins (yolk proteins) while upregulated proteins were enriched in plasma membrane and structural components of muscle cells (Fig. S1B). These results are consistent with previous reports which measured DR-induced proteome changes in mutant backgrounds^7^. The data validate that, within a generation, DR induces substantial proteome-wide changes representing the reduced demand for protein translation under conditions of slower growth.

### Ribosomal proteins are downregulated across generations after DR

The effects of DR on the progeny were significantly weaker than those observed within a generation (Fig. 1C and S1C, mean |log_2_(fold change)| = 0.3 (P0) vs. 0.11 (F1); variance = 0.078 (P0), 0.016 (F1), p = 3.5*10^−126^, KS-test) with only very few proteins significantly down- or upregulated by more than two-fold (Supplemental Table S1b). Overall, protein expression changes in progeny were not, or only very weakly, correlated to those in the parental generation (Fig. S1D, R^2^ = 0.037, p = 1.74*10^−46^). These data suggest that proteome alterations after DR are not globally inherited across generations in *C. elegans*.

Although only few individual proteins passed stringent significance thresholds when analysed individually, the parental diet had a significant overall impact on the proteome composition, as evident from the separation along the first principal component (Fig. S1E, F). This separation suggests that numerous proteins change in expression in response to maternal diet, but since these changes are smaller than two-fold, they may not be detectable at single protein resolution due to measurement noise.

To uncover a potential intergenerational inheritance of the proteome composition of proteins changing less than two-fold, we asked which Gene Ontology (GO) terms representing cellular components were enriched among proteins that were consistently up- or downregulated under DR across generations (|z-score| > 1, Fig. 1D). Nearly all significantly enriched GO terms among the downregulated proteins related to cytoplasmic or mitochondrial ribosomes. The only other GO terms enriched among consistently downregulated proteins referred to large subcellular compartments with little functional specificity. Similarly, upregulated proteins were enriched in only a few GO terms, which related to membrane bound compartments (Fig. 1D, p < 0.01).

For both progeny and parent, nearly all ribosomal proteins were downregulated after DR (139/144 proteins for parents, 139/142 proteins for progeny, Fig. 1E) and their mean expression was significantly reduced (−12%, p = 1.04*10^−22^ for progeny and −31% for parents, p = 4.8*10^−23^, Wilcoxon signed-rank test. Fig. S1G). We did not observe such a dependence on maternal diet for other cytoplasmic proteins. For example, components of the proteasome were nearly unaffected by DR in the progeny (24/40 proteins positive fold-change, 16/40 negative fold-change, 1.5% mean change, p > 0.05, Wilcoxon signed-rank test) although these were strongly biased towards downregulation (41/44 proteins, −23% mean change, p = 1.8*10^−12,^, Wilcoxon signed-rank test) in the parents (Fig. 1E and S1G).

We validated these findings by live imaging using strains carrying endogenously tagged fluorescent proteins sampled by the same procedure as for mass spectrometry. Consistently, ribosomal proteins RPL-34 and RPL-29 maintained reduced expression across generations, while the proteasomal protein RPN-9 was downregulated in mothers but slightly upregulated in their progeny (Fig. 1F). Together, these data reveal that although the proteome is globally altered under DR, the expression of most proteins is reset between generations. However, unlike many other proteins, ribosomal protein downregulation is propagated to the next generation after maternal DR treatment. We note that the quantitative change in ribosome abundance was smaller in progeny than in their parents (Fig. 1F and S1G). This difference nevertheless amounts to a significant change in the total amount of the progeny’s protein mass allocated to ribosomes, given that ribosomal protein comprise an estimated 10% of the worm’s total protein mass^20^.

### Reduced ribosome levels after maternal DR correlates with slow growth and development

To investigate the physiological impact of maternal DR-induced ribosome reduction, we tracked individual progeny development using time-lapse microscopy. We monitored animals carrying an endogenous RPL-34:mCherry tag in agarose-based growth chambers with excess food (*ad libitum*, AL), capturing images every 10 minutes throughout development (Fig. 2A). As previously described^21^, this approach allowed us to derive precise volume growth rates, larval stage durations, as well as the volume at larval molts, which can be readily identified by a halt of growth occurring before cuticular ecdysis^21^ (Fig. 2A).

**Figure 2.**
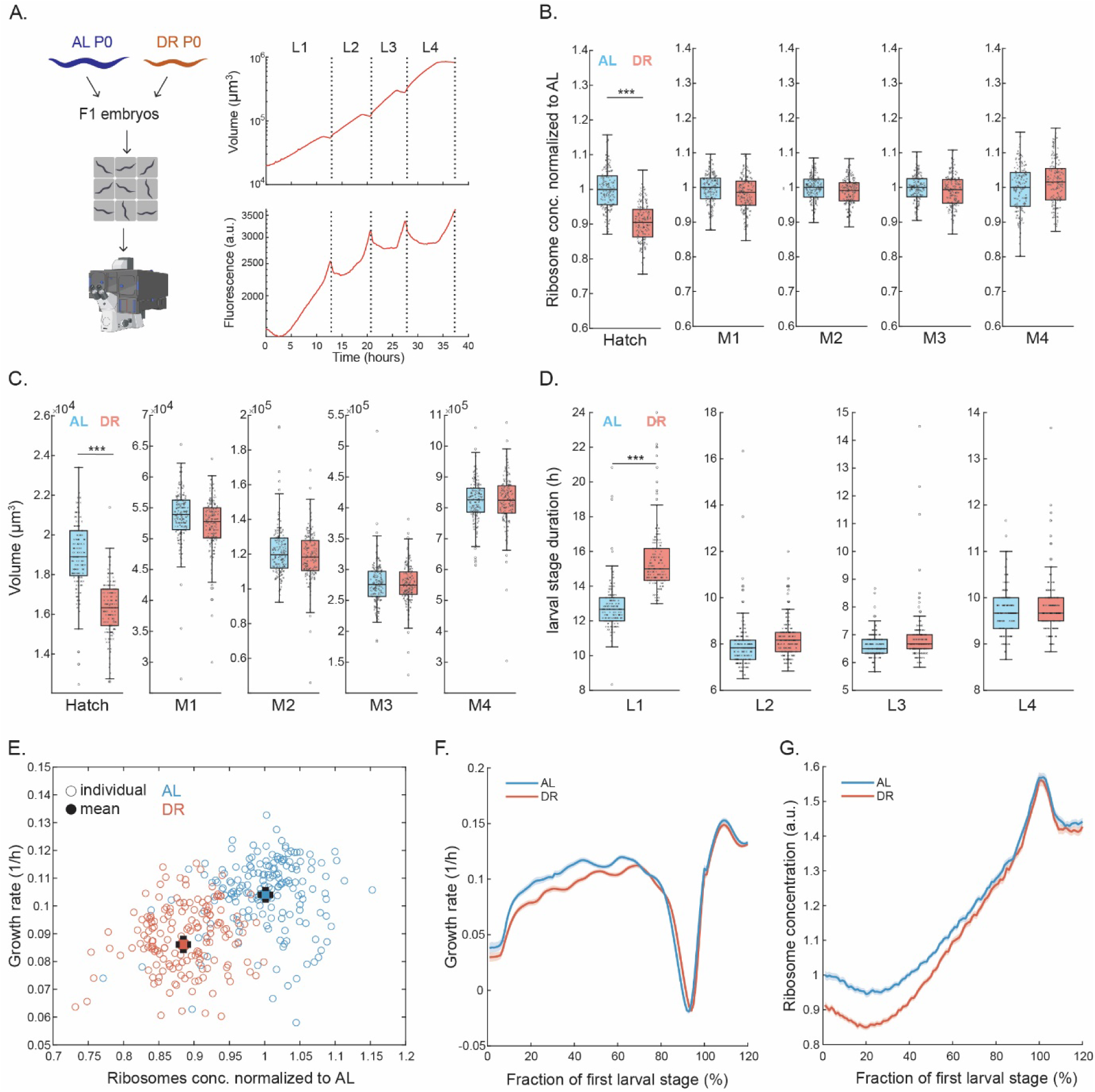
Parental DR reduces ribosome concentration and growth rate of progeny. **(A)** Experimental design and representative trajectory of individual animal growth measurement in agarose chambers. Left: embryonic progeny from AL and DR treated parents were loaded into agarose chambers and monitored throughout their development by live imaging at 10-minute interval. Right: volume and fluorescence trajectory measured for a single individual expressing RPL-34:mCherry over time. Plateaus in volume growth curve represent larval molts and were used to distinguish larval stages (dotted lines). Fluorescence is shown as intensity per pixel. Peaks in fluorescence are due to continued fluorescence production during the growth arrested molt, followed by a rapid expansion after the molt. **(B)** Ribosome concentration (fluorescence per pixel) of progeny of AL-(blue) and DR-(red) treated parents after hatching and at molts M1 to M4. central line: median, box: interquartile range (IQR), whisker: ranges except extreme outliers (>1.5*IQR), individual values: crosses, extreme outliers: circles. For each condition a total of at least n ≥ 170 individuals were measured on at least m ≥ 3 days. *** indicate p<10^−5^ (Wilcoxon rank-sum test). See Supplemental Table S3 for precise sample size and p-values. **(C)** As (B) but for volume after hatching and at ecdysis. **(D)** As (B) but for larval stage durations. **(E)** Correlation between initial growth rate after hatching and ribosome concentrations in fluorescence per pixel at hatching. Individual measurements (circles) and population means (filled circles) shown for offspring AL (blue) and DR (red) treated parents (R² = 0.26, p = 6.69*10^−24^). Error bars: 2*s.e.m. Significance of difference between mean growth rates: p = 2.4*10^−26^ (Wilcoxon rank-sum test) **(F)** Growth rate during L1 development. Individual trajectories were aligned at hatch point and M1 and re-scaled before averaging. Data range between 100% and 120% represents the beginning of L2. Decline in growth rate represents the larval molt. Solid lines: mean, shaded regions: 95% confidence interval. Number of individuals and biological replicates as in (B). **(G)** As (F), but for ribosome concentration (fluorescence per pixel).

Progeny of dietarily restricted mothers (called “DR progeny” from here onwards) had an 11% lower median RPL-34:mCherry intensity than progeny of *ad libitum* fed mothers (called “AL progeny”) (Fig. 2B). These measurements confirm the proteomics and imaging measurements of immobilized animals shown in Fig. 1. As measurements in agarose chambers did not require synchronization at L1 stage, these data also show that the reduction in RPL-34:mCherry occurs independently of larval arrest and is detectable immediately after hatching. In addition to reduced ribosome concentration, DR progeny also had a 13% smaller volume (Fig. 2C). Importantly, however, the observed 11% reduction in RPL-34:mCherry intensity (Fig. 2B, E) was not due to this size decrease, as we measured the change in concentration after size normalization. The total reduction in fluorescence without size normalization was 15%.

Parental DR significantly impacted the speed of early development, delaying the L1 to L2 molt (M1) by an average of 2.7 ± 0.2 hours (s.e.m, p = 1.5*10^−39^, Fig. 2D). This delay results from a combination of the smaller initial hatching volume, and additionally the reduced volume specific growth rate (measured as dV/dt/V = dlog(V)/dt) that was correlated with a reduced ribosome expression (Fig. 2E).

During development, the ribosome concentration progressively increased in DR progeny, reaching levels indistinguishable from that of AL progeny by M1 (Fig. 2B, G). This gradual increase in ribosome levels was paralleled by a matching increase in the growth rate (Fig. 2F), suggesting successful adaptation of DR progeny to AL conditions within the first larval stage. Indeed, beyond M1, development proceeded normally with no significant differences in subsequent larval stage durations, growth rate, ribosome levels, or body volume between DR and AL progeny (Fig. 2B-D).

We conclude that the parental diet impacts the initial ribosome concentration and growth rate after hatching, and that animals restore normal growth rates over several hours of development and in coincidence with adjusting to normal ribosome levels.

### Reduced ribosome concentration after hatch causes slow growth

To determine whether reduced ribosome levels directly caused or were merely correlated with slower growth, we developed a genetic system to quantitatively manipulate ribosome levels at hatch. We inserted an auxin-inducible degron (*aid*) and a GFP tag at the endogenous locus of the *rps-26* ribosomal protein gene and expressed the plant ubiquitin ligase TIR1^22^ under the *sun-1* promoter, which is active in the proximal germline and in the early embryo^23,24^. This approach allowed us to modulate ribosome levels that L1 progeny have at hatch by titrating auxin (indole-3-acetic acid, IAA) concentration in the parental growth medium (Fig. 3A), and use RPS-26:GFP levels to measure ribosomal protein concentrations.

**Figure 3.**
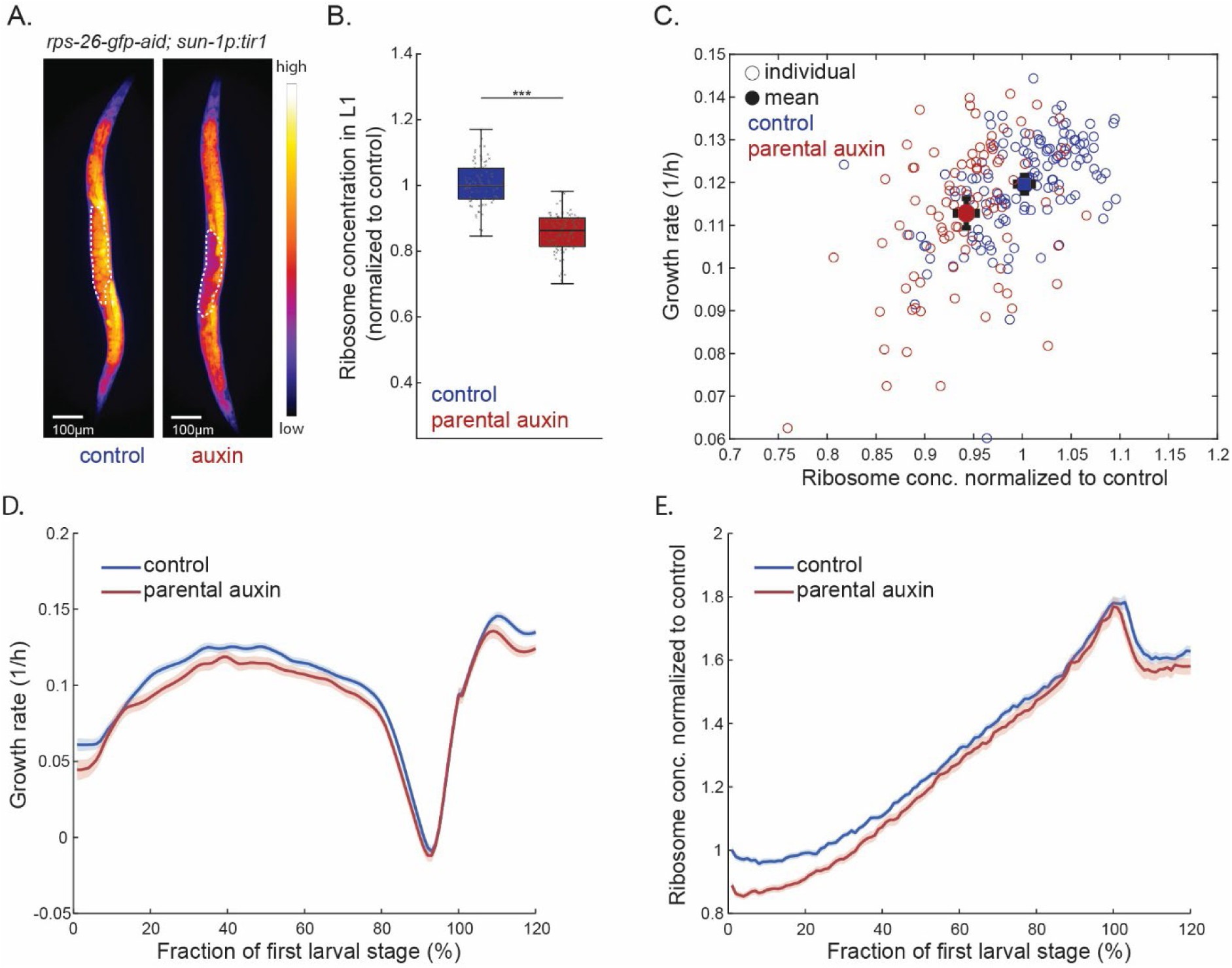
Auxin-induced depletion of RPS-26:GFP in proximal germline and early embryos reduces progeny growth and ribosome levels. **(A)** Fluorescence microscopy image of *C. elegans* with endogenously inserted *rps-26:aid:gfp* tag expressing *sun-1p:tir1* treated with 250 μM auxin (right) or control (left). White dotted line: embryos in uterus. Scale bar: 100 μm. **(B)** Quantification of RPS-26:GFP fluorescence (intensity per pixel) in newly hatched L1 larvae from control and auxin-treated parents. central line: median, box: interquartile range (IQR), whisker: ranges except extreme outliers (>1.5*IQR), individual values: crosses, extreme outliers: circles. Total number of individuals n=131 and 100 for control and auxin-treated, number of experiments m = 3, *** p<10^−33^ (Wilcoxon rank-sum test). **(C)** Correlation between initial growth rates after hatching and ribosome concentrations at hatching. Individual measurements (circles) and population means (filled circles) shown for offspring of control (blue) and auxin-treated (red) parents (R²=0.25, p = 1.3*10^−15^). Error bars: 2*s.e.m. Significance of difference between mean growth rates: p = 1.8*10^−3^ (Wilcoxon rank-sum test). **(D)** Growth rate during L1 development. Individual trajectories were aligned at hatch point and M1 and re-scaled before averaging. Range between 100% and 120% represents the beginning of L2. Solid lines: mean, shaded regions: 95% confidence interval. Number of individuals and biological replicates as in (B). **(E)** As (D), but for Ribosome concentration (fluorescence per pixel).

We established auxin concentrations that, when applied to AL fed mothers, reduced RPS-26:GFP concentration by 14% at hatch (Fig. 3B), close to the reduction observed after parental dietary restriction (Fig. 1). Growth measurements in agarose chambers showed that this modest ribosome reduction was indeed sufficient to recapitulate the slow growth observed after parental DR (Fig. 3C, D). Like progeny of DR animals, progeny depleted for ribosomes by auxin-induced degradation at the time point of hatching similarly reached normal ribosome levels during L1 development and recovered near normal growth rates (Fig. 3D, E). We note that this recovery of ribosome levels occurred slightly more rapidly following auxin-induced experimental depletion than after parental DR (Fig. S2). This difference in the recovery dynamics indicates that while ribosome reduction is sufficient to cause a growth delay, other factors may additionally be involved with the developmental delay caused by parental DR.

### Maternal somatic ribosome levels and progeny ribosome allocation can be uncoupled

We next asked if reduced maternal growth rates and ribosome levels inherently impact the ribosome concentration in the progeny, or if ribosome levels in the maternal soma and in the progeny can be uncoupled. We used a strain expressing Tir1 under the *eft-3* promoter, a transgene that is active in all somatic cells, but silenced in the germline, and an endogenously tagged *rpl-22:aid* ribosomal protein to selectively deplete ribosomes in the parental soma. Additionally, we fluorescently tagged another ribosomal protein (RPL-34:mCherry) for the measurement of ribosomal protein concentrations. Addition of auxin early in development led to strong soma-specific depletion of ribosomes and caused developmental arrest and sterility (Fig. S3A, B). To ask how somatic ribosome depletion in mothers impacts the next generation, we applied auxin from the fourth larval stage onwards, which did not abolish reproduction, but nevertheless reduced maternal growth and ribosome concentration (Fig. 4A, B). Despite this significant reduction in maternal growth upon addition of auxin, ribosome levels and growth rates were nearly unaffected in L1 larvae of the next generation (Fig. 4C, D), suggesting that ribosome levels in the progeny can be uncoupled from those in the parental soma.

**Figure 4.**
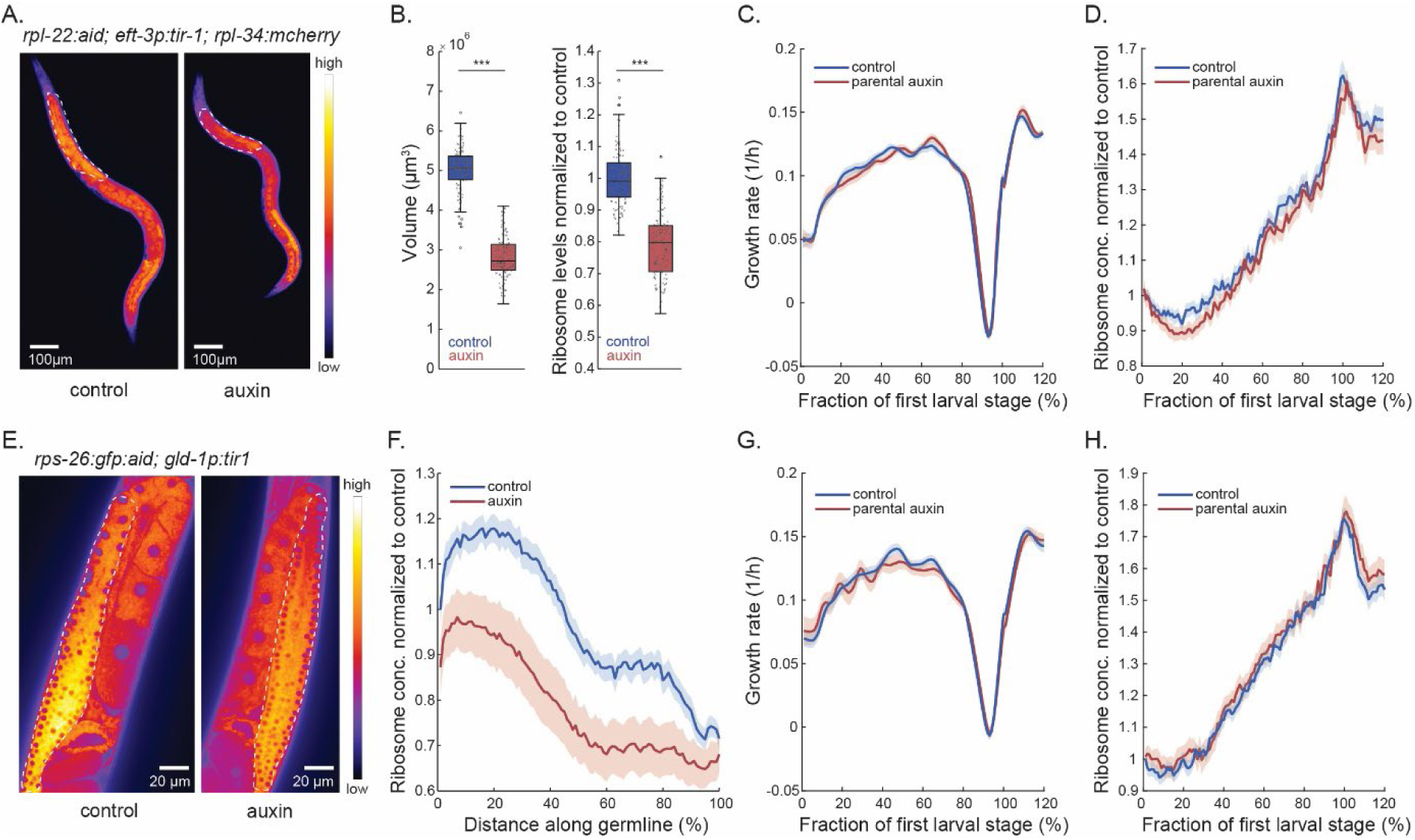
Progeny ribosome levels are robust to parental somatic as well as distal germline ribosome depletion. **(A)** Fluorescence microscopy image of *C. elegans* with *rpl-22:aid* and endogenously inserted *rpl-34:mcherry* tag expressing *eft-3p:tir1* treated with 500 μM auxin (right) or control (left). White dotted line: ribosomes depleted in intestine. Scale bar: 100 μm. **(B)** Quantification of volume and RPL-34:mCherry fluorescence (intensity per pixel) in control and auxin-treated individuals. central line: median, box: interquartile range (IQR), whisker: ranges except extreme outliers (>1.5*IQR), individual values: crosses, extreme outliers: circles. Total number of individuals n=116 and 119 for control and auxin-treated, number of experiments m = 2, *** p<10^−30^ (Wilcoxon rank-sum test). **(C)** Growth rate of F1 progeny of *rpl-22:aid; eft-3p:tir1; rpl-34:mcherry* animals after parental auxin treatment. Individual trajectories were aligned at hatch point and M1 and re-scaled before averaging. Range between 100% and 120% represents the beginning of L2. Solid lines: mean, shaded regions: 95% confidence interval. n = 82 and 63 individuals from m = 2 experiments. **(D)** As (C), but for RPL-34:mCherry concentration (fluorescence per pixel). **(E)** RPS-26:GFP fluorescence of *rps-26:gfp:aid;glp-1p:tir1* auxin-treated (right) and untreated (left). White dotted line: ribosome depletion in distal germline. **(F)** Spatial distribution of RPS-26:GFP intensity (fluorescence per pixel) along the germline from distal tip (0%) to most proximal oocyte (100%). Blue: control, red: 1000 μM auxin. n = 55 and 55 individuals from m = 3 experiments. Lines: population mean ± 95% CI. **(G)** Growth rate of progeny of *rps-26:gfp:aid;glp-1p:tir1* animals after parental auxin treatment. Individual trajectories were aligned at hatch point and M1 and re-scaled before averaging. Range between 100% and 120% represents the beginning of L2. Solid lines: mean, shaded regions: 95% confidence interval. n = 108 and 79 individuals from m = 2 experiments. **(H)** As (G), but for RPS-26:GFP concentration (fluorescence per pixel).

To further validate that progeny and parental ribosome levels can be uncoupled, we selectively depleted RPS-26 in the parental germline using Tir1 expressed under the *gld-1* promoter. This promoter is specifically expressed in the distal portion of the germline, but not active in the proximal germline where oocytes are formed, nor in embryos^25,26^. Consistently, auxin treatment depleted ribosome levels in the distal region of the germline (Fig. 4E). However, again, ribosome levels and growth rate in the progeny were unchanged (Fig. 4G, H). These findings reveal a remarkable robustness of progeny ribosome allocation to perturbations of ribosomes in parental tissues, suggesting that mechanisms controlling progeny ribosomes do not operate immediately downstream of parental somatic ribosome levels or somatic growth rates *per se*.

### Somatic insulin-like signalling does not impact progeny ribosomes

Insulin and insulin-like growth factor signalling (IIS) through the receptor *daf-2* and its downstream effector *daf-16* play a central role in many DR-induced physiological changes, including the regulation of progeny size and starvation resistance^14,18,27^. To ask if reduced IIS also reduces progeny ribosome control and growth, we employed an available *daf-2-aid* allele^28^ with a soma-specific *eft-3p:tir1* transgene and measured RPL-34:mCherry in progeny animals.

Since even low auxin dose triggered highly penetrant developmental arrest as *dauer* larvae^28^, we exposed animals to auxin from the L4 stage onwards and assessed progeny outcomes. Auxin-induced DAF-2 depletion increased progeny size by 17% (Fig. 5A), as was previously reported for genetic alleles of *daf-2(e1370)* and maternal somatic *daf-2* RNAi^14^, showing the effectiveness of DAF-2 depletion. However, DAF-2 depletion did not impact the progeny’s ribosome concentration normalized to size (Fig. 5B, D) or its volume specific growth rate (Fig. 5C), suggesting that somatic IIS does not mediate intergenerational ribosome control.

**Figure 5.**
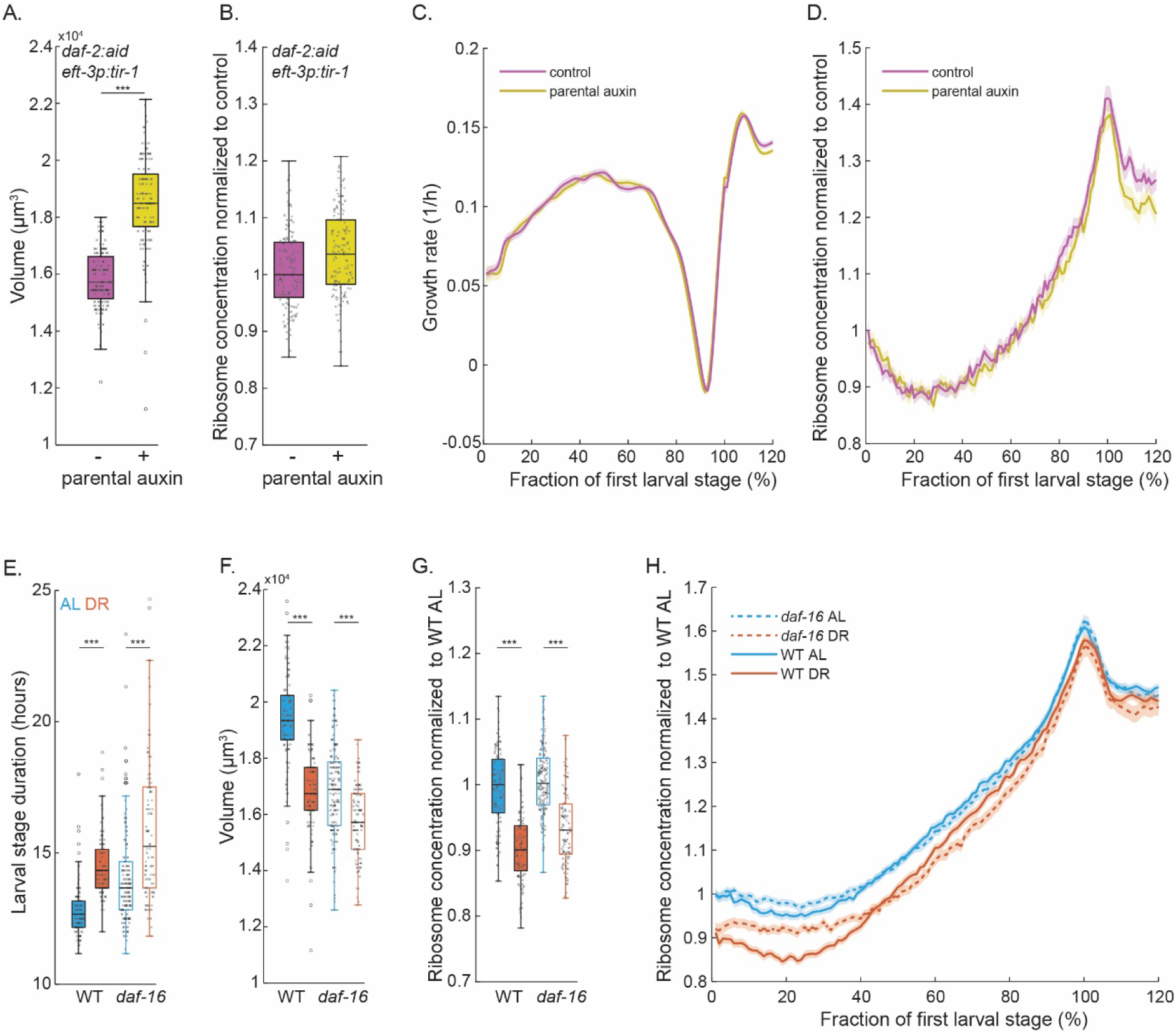
Insulin/Insulin-like Signalling (IIS) is not involved in intergenerational ribosome control. **(A)** Volume of L1 progeny at hatch from parents somatically depleted for DAF-2-AID with 500 μM auxin (yellow) and control (magenta). central line: median, box: interquartile range (IQR), whisker: ranges except extreme outliers (>1.5*IQR), individual values: crosses, extreme outliers: circles. Number of individuals n = 138 and 141 from m = 2 experiments. p = 9*10^−37^ (Wilcoxon rank-sum test). **(B)** Same as (A), but for RPL-34:mCherry concentration (pixel intensity normalized to control). Parental auxin does not reduce progeny ribosome concentration but slightly increases it (p = 0.001, Wilcoxon rank-sum test). **(C)** Growth rate of progeny during L1 development for parental depletion of DAF-2 and control. Individual trajectories were aligned at hatch point and M1 and re-scaled before averaging. Range between 100% and 120% represents the beginning of L2. Solid lines: mean, shaded regions: 95% confidence interval. Number of individuals and biological replicates as in (A). **(D)** As (C), but for RPL-34:mCherry concentration (fluorescence per pixel). **(E)** Same as (A), but for L1 larval stage duration and for parental AL and DR in wild type (filled boxes) and *daf-16(mu86)* mutants (open boxes). For each condition a total of at least n ≥ 100 individuals were measured on at least m ≥ 3 days. *** indicate p<10^−5^ (Wilcoxon rank-sum test). See Supplemental Table S4 for precise sample size and p-values. **(F)** Same as (E), but for volume at hatching. **(G)** Same as (E), but for RPL-34:mCherry concentration at hatch (fluorescence per pixel) normalized to wild type AL. **(H)** RPL-34:mCherry concentration as described for (C), but for indicated genotypes and parental dietary treatments.

To corroborate these findings, we investigated the role of *daf-16* in intergenerational ribosome control using its null allele *daf-16(mu86)*. These mutants showed profound sensitivity to parental DR, exhibiting a substantial growth delay (Fig. 5E) and a smaller size after hatching (Fig. 5F) when measured in agarose chambers. These data thereby confirm the role of IIS in responding to DR within and across generations. However, as was the case for parental *daf-2* depletion, mutation of *daf-16* in parents and progeny and did not affect RPL-34:mCherry concentrations at hatching or its response to parental DR (Fig. 5G), indicating that developmental delay of the *daf-16(mu86)* mutant was not caused by changes in ribosome levels. While ribosome levels at hatch were unaffected by *daf-16* mutation, the expression dynamics during L1 growth were slightly altered, consistent with a role of *daf-16* in ribosome control during somatic growth, but not across generations (Fig. 5H).

### The mTORC1 pathway regulator RAGA-1 intergenerationally controls ribosomes

In addition to IIS, the mTORC1 signalling pathway represents a critical regulator of organismal growth across eukaryotes, promoting translation and ribosome synthesis under nutrient rich conditions and activating autophagy under starvation^29–31^. To investigate the role of mTORC1 in intergenerational ribosome control, we depleted RAGA-1, the sole *C. elegans* ortholog of the mTORC1 activators RagA/B^32–34^, in the soma using AID with an *eft-3p:tir1* transgene (Fig. 6A).

**Figure 6.**
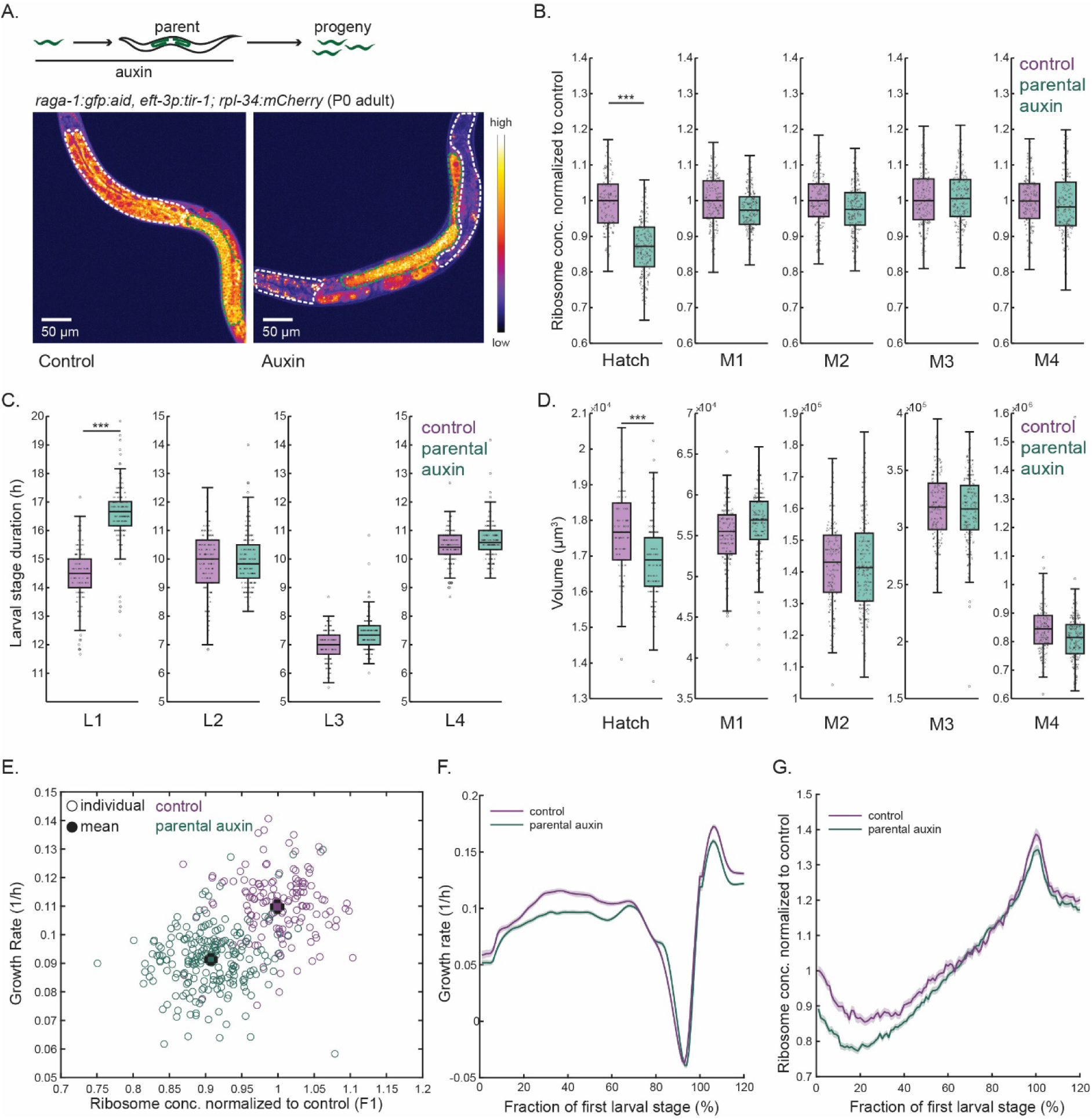
Depletion of RAGA-1 in parental soma phenocopies parental dietary restriction. **(A)** Top: Experimental design. RAGA-1 was parentally depleted in the soma using a somatically expressed *eft-3p:tir1* transgene and an endogenously inserted *raga-1-gfp-aid* allele. 500 μM auxin was applied to parental generation. Bottom: RAGA-1:GFP fluorescence of *raga-1:gfp:aid; eft-3p:tir1; rpl-34:mCherry* auxin-treated (right) and untreated (left). White dashed outline: intestine (soma), green dotted outline: germline. Scale bar: 50 μm. **(B)** RPL-34:mCherry concentration (fluorescence per pixel) after parental somatic RAGA-1 depletion (green) and control (magenta) normalized to control. central line: median, box: interquartile range (IQR), whisker: ranges except extreme outliers (>1.5*IQR), individual values: crosses, extreme outliers: circles. For each condition a total of at least n = 150 individuals were measured on at least m = 3 days. *** indicate p<10^−5^ (Wilcoxon rank-sum test). See Supplemental Table S5 for precise sample size and p-values. **(C)** As (B), but for larval stage duration. **(D)** As (B) but for volume after hatching and at larval molts. **(E)** Correlation between initial growth rate after hatching and ribosome concentrations at hatching. Individual measurements (circles) and population means (filled circles) shown for control and (magenta) and after parental RAGA-1 depletion (cyan). (R²=0.25, p = 2.3*10^−24^). Error bars: 2*s.e.m. Significance of difference between mean growth rates: p = 2*10^−35^ (Wilcoxon rank-sum test). **(F)** Growth rate of progeny with and without parental RAGA-1 depletion. Individual trajectories were aligned at hatch point and M1 and re-scaled before averaging. Range between 100% and 120% represents the beginning of L2. Solid lines: mean, shaded regions: 95% confidence interval. Number of individuals and biological replicates as in (B). **(G)** As (F), but for RPL-34:mCherry concentration (intensity per pixel).

Consistent with our previous findings^34^ and a role of mTORC1 in ribosome biogenesis, RAGA-1 depletion slowed-down organismal development (Fig. S4A) and reduced the expression of RPL-34:mCherry (Fig. S4B). Critically, however, and unlike somatic depletion of DAF-2 (Fig. 5) or depletion of ribosomes in the distal germline (Fig. 4), the somatic depletion of RAGA-1 also reduced RPL-34:mCherry concentrations in the progeny (Fig. 6B). Parental RAGA-1 AID also phenocopied other parental DR-associated characteristics, slowing down development during the L1 stage (Fig. 6C), reducing the size of L1 larvae at hatching (Fig. 6D), and slowing down the growth rate after hatching in correlation with reduced ribosome levels (Fig. 6E-G). As for parental DR, these phenotypic changes converged towards those of untreated animals within the first larval stage (Fig. 6F, G), such that animals experiencing parental RAGA-1 depletion were indistinguishable from untreated controls from the L2 stage onwards (Fig. 6B-D).

In summary, somatically reduced parental RAGA-1 comprehensively recapitulates the intergenerational phenotypic plasticity induced by dietary restriction. These data suggest that mTORC1 signalling in the parental soma influences the ribosome concentration in the progeny and thereby contributes to growth control across the soma-germline barrier that separates two generations.

## Discussion

Organisms adjust their proteome composition to their environmental conditions. Previous work has systematically analysed this response in unicellular microbes and revealed that many proteins scale linearly with the growth rate^1,2^. A similar relation occurs in animal cells for ribosomes, which scale with the growth rate and the anabolic demand of cells^7^. Here, we used dietary restriction of *C. elegans* as a model to investigate an aspect of this control specific to multicellular life: proteome regulation across generations in response to maternal diet.

A key finding of this study is that most diet-induced changes to the proteome are not propagated across generations in *C. elegans*, highlighting a marked difference between unicellular and multicellular organisms. However, ribosomes are an exception to this rule and are downregulated in the dietarily restricted parental generation, as well as in their progeny (Fig. 1D).

The first 15 hours of *C. elegans* embryonic development take place inside the eggshell, which restricts exchange with environmental conditions. The proteome composition at hatching is therefore mainly determined by material and/or signalling molecules deposited by the mother prior to eggshell formation. Our observation of reduced ribosome levels in the progeny of dietary restricted animals suggests a mechanism in which mothers adjust ribosome provisioning according to the anticipated need of their offspring. Notably, the reduction in progeny ribosome levels (11%) is less strong than in dietary-restricted parents (∼30%). This strategy may balance potential metabolic savings of reduced ribosome production with being prepared for a potentially improved environment at the time of hatching.

Reduced ribosome levels upon maternal DR have significant physiological consequences, slowing down progeny growth when feeding *ad libitum* until normal ribosome concentrations are restored (Fig. 2). Targeted degradation of ribosomes in hatching larvae by a similar amount recapitulated this slower growth rate, showing that ribosomes are indeed limiting for growth after maternal DR. While these data reveal ribosomes as a rate-limiting bottleneck in intergenerational growth control, they do not exclude that additional, ribosome-independent, mechanisms are also involved in intergenerational inheritance upon maternal DR.

Reduced ribosome levels in progeny cause a significant growth disadvantage when conditions improve from DR to AL across generations. This disadvantage raises the question of what benefit lower ribosome levels in the progeny may have. At least three non-mutually exclusive explanations are possible. First, the reduced allocation of resources to ribosomes may allow for improved growth under continued dietary restriction. Testing this possibility may become possible in the future with further technological developments to precisely titrate food concentration in experimental systems suitable for long term imaging at scale. Second, performance of progeny in other tasks, such as resistance to stress or faster growth could be improved by reduced ribosomes. Indeed, DR progeny have improved robustness of germline development to prolonged starvation^27^, although it is not certain that these benefits are directly related to ribosome expression. Third, reduced progeny ribosomes could accelerate the reproduction rate of the mother, i.e. the rate of eggs produced per time, as fewer resources need to be spent on ribosome production.

Although lower ribosome levels may yield progeny of reduced quality if they meet conditions with high food availability, such a reduction is not expected to negatively impact progeny growth and fitness if dietary restriction continues across generations. Ultimately, the optimal allocation of ribosomes to progeny may depend on how well the environment experienced by the mother is predictive for what the progeny will encounter, i.e. on how likely the environmental conditions are to change within the 15 hours between eggshell formation and hatching.

Previous studies have documented various cases where the transcriptional or translational control of specific genes is maternally influenced through chromatin modifications or small RNAs^10,35,36^, but have not addressed intergenerational effects on core metabolic machinery. An important open question is the exact chain of signalling events that controls ribosomes across generations. The majority of ribosomes present in L1 larvae at hatch is provided by maternal synthesis^37^. The intergenerational control of ribosomes may thus not require such deposition of regulatory molecules but be determined directly by the number of ribosomes deposited in the oocyte. Depletion of ribosomes in the maternal soma did not impact progeny ribosome levels and depletion in the distal germline did not impact ribosomes in the progeny or the proximal germline (Fig. 4). This robustness to ribosome depletion indicates that most offspring ribosomes are synthesised in the proximal germline immediately prior to or during oogenesis. Our results using depletion of RAGA-1 suggest that parental mTORC1 signalling plays an important role in controlling the ribosome content of the progeny. Importantly, since we depleted RAGA-1 specifically in the maternal soma, regulation of ribosomes in progeny likely involves cell communication across the soma-germline boundary, opening exciting opportunities for mechanistic dissection of this control.

In conclusion, our work provides a systematic study of intergenerational proteome control in response to maternal diet. Our work highlights the allocation of proteome resources to ribosomes as a new mechanism for the intergenerational transmission of phenotypic plasticity that may prepare progeny for environmental conditions that they are most likely to encounter.

## Methods

### *C. elegans* strains and maintenance

Animals were grown and maintained at 25°C using OP50-1 *E. coli* on nematode growth medium (NGM) according to standard procedures. The following strains were used in this study:

N2: Bristol N2 wild isolate: used in Fig. 1A-E, Fig. S1
PHX1880: *rpl-34(syb1880) [rpl-34:RPL-34:mCherry] IV* used in Fig. 1E, Fig. 2
PHX1945: *rpl-29(syb1945) [rpl-29:GFP]* used in Fig. 1E
CER620: *ubh-4(cer68[ubh-4::eGFP]) rpn-9(cer203[rpn-9::wrmScarlet]) II* used in Fig. 1E
wBT340: *rubSi4[rps-26::aid::gfp] I; unc-119(ed3) III; ieSi38 [sun-1p::TIR1::mRuby::sun-1 3’UTR + Cbr-unc-119(+)] IV* used in Fig. 3
wBT271: *rpl-34(syb1880) [rpl-34:RPL-34:mCherry] IV, ieSi57 [eft-3p::TIR1::unc-54 3’UTR + Cbr-unc-119(+)] II (1.73). rpl-22(ohm14)[rpl-22::AID::3xFLAG(codon optimized)] II (−2.58)* used in Fig. 4A-D, Fig. S3A,B
SUR21: *rubSi4[rps-26::aid::gfp] I ; [Pgld-1::TIR1::mRuby::gld-1 3’UTR] II* used in Fig. 4E-H
wBT440: *ieSi57 II; daf-2(bch40) III; rpl-34(syb1880) [rpl-34:RPL-34:mCherry] IV:5.83* used in Fig. 5A-D
wBT371: *daf-16(mu86)I; rpl-34(syb1880) [rpl-34:RPL-34:mCherry] IV:5.83* used in Fig. 5E-H
wBT316: *raga-1(wbm40) [raga-1::AID::EmGFP] II; xeSi376[Peft-3::TIR1::mRuby::unc-54 3’UTR, cb-unc-119(+)] III; rpl-34(syb1880) [rpl-34:RPL-34:mCherry]* used in Fig. 6, Fig. S4A,B

### Liquid culture growth medium for dietary restriction

DR was attained by growth in liquid culture. For liquid culture, the bacterial strain *Escherichia coli* HB101 was diluted in S-basal (supplemented with 1M Potassium citrate, 1M CaCl_2_, 1M MgSO_4_, trace metal solution, 5mg/l cholesterol, 100mg/l carbenicillin, 10mg/l Nystatin)^38^ as a food source at 2*10^8^ cfu/ml for DR conditions. AL conditions were identical, except with 10^9^ cfu/mL. Prior to liquid culture, animals were grown on NGM plates at 25°C for at least three generations. To obtain synchronized populations, gravid adults were bleached, and their embryos were allowed to hatch overnight in S-Basal buffer without ethanol or cholesterol. These synchronized L1 larvae were then cultured in liquid culture medium a density of 100 worms/ml in 50-ml falcon tubes rotated at 25°C. At this worm density no change in food concentration due to consumption was detectable during the course of our experiment.

### Mass spectrometry sample preparation

For mass spectrometry of adults, animals subjected to AL feeding or DR were collected after cultivation in liquid for 3 or 5 days, respectively, starting from synchronized L1 larvae. For F1 progeny, eggs were released from gravid adults cultured under DR or AL by bleaching and remnants of incompletely bleached adults were removed by filtering through a 40 μm mesh (Pluriselect). L1s were allowed to hatch overnight in S-Basal (without cholesterol or ethanol) on an orbital rotor and unhatched eggs were removed by filtering through a 20 μm mesh (Pluriselect) to obtain synchronized, arrested L1 larvae. For each condition three biological replicates were obtained. Number of animals per repeat: 5’000, 10’000, and 100’000 animals each for AL adults, DR adults, and L1 progeny, yielding 30 – 50 μg protein for each sample. To extract protein, worm pellets were resuspended in 100 µL of lysis buffer (PBS pH7.4 containing 0.025% Triton X-100 and protease inhibitors (Sigma P8340)) and flash frozen in liquid nitrogen followed by thawing at 37°C three times and subsequently sonicated for 15 minutes using 30-second on/off cycles in a sonicator bath (Bioruptor). Insoluble protein was removed by centrifugation and the concentration of soluble protein was determined by a micro-BCA assay (Thermo Fischer Scientific).

### TMT Mass spectrometry

The raw output files of FragPipe (protein.tsv files) were processed using the R programming language (ISBN 3-900051-07-0). Contaminants and reverse proteins were filtered out and only proteins that were quantified with at least 2 razor peptides (Razor.Peptides >= 2) were considered for the analysis. 7475 proteins passed the quality control filters. Log_2_ transformed raw TMT reporter ion intensities (’channel’ columns) were first cleaned for batch effects using the ‘removeBatchEffect’ function of the limma package^39^ and further normalized using the ‘normalizeVSN’ function of the limma package (VSN - variance stabilization normalization^40^). Proteins were tested for differential expression using a moderated t-test by applying the limma package (’lmFit’ and ‘eBayes’ functions). The replicate information was added as a factor in the design matrix given as an argument to the ‘lmFit’ function of limma. A protein was annotated as a hit with a false discovery rate (FDR) smaller 0.05 and an absolute fold-change of greater 2.

### Single time-point imaging

*C. elegans* strains with endogenously tagged ribosomes or proteasomes, collected at early adulthood stage or as overnight hatch-synchronized larvae were imaged at single time point. Worms were mounted onto 2% agarose pads on slides and a drop of 10mM levamisole was used to anesthetize the worms. For single time point imaging in Fig. 4C and 6A, a Nikon Ti2 spinning disk microscope (CREST optics v3) was used with a 60x oil objective (N = 1.4). Single time point imaging in Fig. 1 was conducted on Nikon Ti2 epifluorescence microscope using a 10x air objective (NA =0.45) for adults and using a 20x air objective (NA =0.75) for L1 larvae.

### Live imaging in microchambers

For live imaging of progeny of animals cultured in liquid by DR or AL, adults were collected from liquid culture filtered through a 40 µm mesh cell strainer (Pluriselect) to remove younger animals. Egg laying was then induced by incubating adults in 35 mM serotonin in S-Basal on an orbital rotor at 25°C for 1 hour and the released embryos were used for loading into agarose-based growth chambers. For experiments using parental auxin treatment (Fig. 3, 4, 5A-D, 6) the parental generation was cultivated on standard NGM plates supplemented with auxin and eggs were picked directly from these plates for loading into agarose chambers.

Arrayed agarose microchambers were manufactured using 4.5% agarose dissolved in S-Basal (containing 5 μg/ml cholesterol). An inverse replicate was created from a polydimethylsiloxane (PDMS) stamp, as described by Turek et al. 2014^42^ with minor modifications described by Stojanovski et al.^21^. Chamber dimensions were 600 μm x 600 μm x 20 μm. For imaging larval development, chambers were filled with the bacterial strain OP50-1 scraped from a standard NGM plate as a food source, 1.5 to 2 fold stage embryos were manually placed in individual chambers using an eyelash pick, and agarose chambers were sealed by inverting onto a 3.5cm wide dish with a high optical quality gas-permeable polymer bottom (ibidi). 3% low melting temperature agarose dissolved in S-basal (containing 5 μg/ml cholesterol) was pipetted around the agarose chamber array and topped with ∼300-500 μl PDMS to prevent evaporation. Finally, the dish was sealed with parafilm and mounted on a custom-made sample holder for microscopy.

All time lapse imaging was conducted on a Nikon Ti2 epifluorescence microscope equipped with a Hamamatsu Flash 4 sCMOS camera using a 10x air objective (NA =0.45). Software-based autofocus was performed at each time point using the default parameters in Nikon NIS software. GFP excitation was at 470 nm, and mCherry excitation was at 575 nm, with 10 ms exposure times using a SpectraX light source (Lumencor). Images were acquired every 10 minutes until worms reached adulthood. Growth, development, and fertility of worms were unaffected by these illumination conditions.

### Image analysis

Worm segmentation and fluorescence measurements were conducted as previously described using a custom made computational pipeline in Matlab^21^. In brief, the outline of the animals was detected using the Sobel algorithm by MATLAB edge() function, followed by connecting the nearest endpoints to close gaps in the detected contours. Segmented worms were then straightened computationally, and the volume was inferred assuming rotational symmetry. Images where worm segmentation and/or straightening was faulty were detected by a decision-tree-based classifier based on shape features of the straightened animals. An ensemble of 20 bagged decision trees was trained on a subset of manually annotated images using MATLAB’s TreeBagger() function. Images classified as faulty straightening or segmentation were excluded from further analysis.

### Computation of volume and growth rates

The volume of individual worms was calculated at each time point from binary masks, assuming rotational symmetry. Larval stage transitions were identified by detecting the maximum of the second time derivative of the logarithm of animal volume, followed by manual correction using a MATLAB graphical user interface. Larval volumes at each molt (M1 to M4) were calculated using a linear regression of the volumes from the ten time points before the molt. For the volume at birth, regression was based on the ten time points after hatching. To compare growth rate and fluorescence trajectories of strain and conditions with different larval stage durations, each individual’s trajectory was scaled by linearly interpolating each larval stage into 100 evenly spaced points, averaging across all individuals, and plotting as a function of larval stage progression^21^. Growth rates were calculated from the worm volumes, which were median-filtered with a three-time-point window and further smoothed over 15 time points using MATLAB’s smooth() function with the “rlowess” option. Growth rates were normalized to the duration of each larval stage, and individual signals were interpolated into 100 points per stage.

Growth rates and ribosome concentrations were analysed in individual animals using a 3.7-hour measurement window centred at 4.3 hours after hatching. Growth rates were computed by performing a robust linear regression on the natural logarithm of volume over this window, after applying a 3-point median filter. The slope of this regression represents the instantaneous relative growth rate (dlog(V)/dt). For ribosome concentration, raw fluorescence intensities were background-subtracted, normalized to the animal’s area, and then median-filtered using a 3-point window. The resulting values were normalized to the mean fluorescence of control conditions to obtain relative ribosome concentrations. Data points where either growth rates or normalized ribosome levels deviated by more than 3 standard deviations from their respective means were excluded as outliers.

### Quantification of germline lengths and fluorescence intensity

Strains expressing RPS-26:GFP:AID and TIR1 under *gld-1* promoter were cultured at 25°C on 2% NGM plates until the L4 stage. Worms were then transferred to either auxin-containing plates (1mM) or control plates containing ethanol until adulthood.

Confocal z-stacks were obtained from adult animals expressing RPS-26:GFP:AID. The midline of each germline was manually traced using FIJI in the central focal plane of the germline from the distal to the spermatheca. For each individual and gonad, a line profile of the fluorescence intensity with the width adjusted to match the width of the gonad. To account for variations in germline length each trace was rescaled to the same length followed by interpolation at 100 points equally spaced points prior to averaging individual traces from the same treatment group.

## Supporting information

Supplemental Tabls S1a

## Acknowledgements

We are thankful to Cihan Elci for technical assistance, Suzan Ruijtenberg (Utrecht University) for sharing strains prior to publication, and Frank Stein and Jennifer Schwarz (EMBL Heidelberg) for Proteomic analysis. We acknowledge support by the Microscopy Imaging Center at the University of Bern. This work received funding from the Swiss National Science Foundation (SNSF) in the form of an Eccellenza Professorial Fellowship (PCEFP3_181204) to B.D.T., the Novartis Foundation for Medical-Biological Research (Grant #20A011), and the Berne University Research Foundation. B.D.T. is thankful to EMBO young investigator program (#5623) for support. N.S. received support from European Research Council under the European Union’s Horizon 2020 Research and Innovation Programme (grant agreement no. 852201), The Spanish Ministry of Economy, Industry and Competitiveness to the EMBL partnership, the Centro de Excelencia Severo Ochoa (CEX2020-001049-S, MCIN/AEI/10.13039/501100011033), the CERCA Programme/Generalitat de Catalunya (NS),The Spanish Ministry of Economy, Industry and Competitiveness Excelencia awards PID2020-115189GB-I00 and PID2023-147692NB-I00. Some strains were provided by the CGC, which is funded by NIH Office of Research Infrastructure Programs (P40 OD010440).

## Author contributions

B.D.T and S.P. conceived the study, wrote the manuscript. B.D.T and S.P. edited the manuscript. S.P. and K.S. conducted all other experiments. S.P., B.D.T. and J.T. performed computational and statistical analyses. S.P., B.D.T. and N.S. created and validated strains used in the study.

## References

1. Scott, M., Gunderson, C. W., Mateescu, E. M., Zhang, Z. & Hwa, T. Interdependence of Cell Growth and Gene Expression: Origins and Consequences. Science 330, 1099–1102 (2010).

2. Schaechter, M., MaalOe, O. & Kjeldgaard, N. O. Dependency on Medium and Temperature of Cell Size and Chemical Composition during Balanced Growth of Salmonella typhimurium. Journal of General Microbiology 19, 592–606 (1958).

3. Scott, M., Klumpp, S., Mateescu, E. M. & Hwa, T. Emergence of robust growth laws from optimal regulation of ribosome synthesis. Molecular Systems Biology 10, 747 (2014).

4. Towbin, B. D. et al. Optimality and sub-optimality in a bacterial growth law. Nat Commun 8, 14123 (2017).

5. Kim, M.-S. et al. A draft map of the human proteome. Nature 509, 575–581 (2014).

6. Jiang, L. et al. A Quantitative Proteome Map of the Human Body. Cell 183, 269–283.e19 (2020).

7. Depuydt, G. et al. Reduced insulin/insulin-like growth factor-1 signaling and dietary restriction inhibit translation but preserve muscle mass in Caenorhabditis elegans. Mol Cell Proteomics 12, 3624–3639 (2013).

8. Weismann, A. The Germ-Plasm: A Theory of Heredity. Translated by W. Newton Parker and Harriet Rönnfeldt. (Scribner, New York, 1893). doi:10.5962/bhl.title.25196.

9. Bonduriansky, R. & Day, T. Nongenetic Inheritance and Its Evolutionary Implications. Annu. Rev. Ecol. Evol. Syst. 40, 103–125 (2009).

10. Perez, M. F. & Lehner, B. Intergenerational and transgenerational epigenetic inheritance in animals. Nat Cell Biol 21, 143–151 (2019).

11. Agrawal, A. A., Laforsch, C. & Tollrian, R. Transgenerational induction of defences in animals and plants. Nature 401, 60–63 (1999).

12. Burton, N. O. et al. Insulin-like signalling to the maternal germline controls progeny response to osmotic stress. Nat Cell Biol 19, 252–257 (2017).

13. Jenkins, N. L. & Hoffmann, A. A. Genetic and maternal variation for heat resistance in Drosophila from the field. Genetics 137, 783–789 (1994).

14. Hibshman, J. D., Hung, A. & Baugh, L. R. Maternal Diet and Insulin-Like Signaling Control Intergenerational Plasticity of Progeny Size and Starvation Resistance. PLoS Genet 12, e1006396 (2016).

15. Bošković, A. & Rando, O. J. Transgenerational Epigenetic Inheritance. Annu. Rev. Genet. 52, 21–41 (2018).

16. Skvortsova, K., Iovino, N. & Bogdanović, O. Functions and mechanisms of epigenetic inheritance in animals. Nat Rev Mol Cell Biol 19, 774–790 (2018).

17. Henderson, I. R. & Jacobsen, S. E. Epigenetic inheritance in plants. Nature 447, 418–424 (2007).

18. Mata-Cabana, A., Pérez-Nieto, C. & Olmedo, M. Nutritional control of postembryonic development progression and arrest in Caenorhabditis elegans. in Advances in Genetics vol. 107 33–87 (Elsevier, 2021).

19. Baugh, L. R. & Hu, P. J. Starvation Responses Throughout the Caenorhabditiselegans Life Cycle. Genetics 216, 837–878 (2020).

20. Tuomaala, J. et al. Selective autophagy of ribosomes balances a tradeoff between starvation survival and growth resumption. Preprint at 10.1101/2024.08.28.609383 (2024).

21. Stojanovski, K., Großhans, H. & Towbin, B. D. Coupling of growth rate and developmental tempo reduces body size heterogeneity in C. elegans. Nat Commun 13, 3132 (2022).

22. Sepers, J. J., et al. The mIAA7 degron improves auxin-mediated degradation in *Caenorhabditis elegans*. G3 Genes|Genomes|Genetics 12, jkac222 (2022).

23. Ashley, G. E. et al. An expanded auxin-inducible degron toolkit for *Caenorhabditis elegans*. Genetics 217, iyab006 (2021).

24. Zhang, L., Ward, J. D., Cheng, Z. & Dernburg, A. F. The auxin-inducible degradation (AID) system enables versatile conditional protein depletion in *C. elegans*. Development dev.129635 (2015) doi:10.1242/dev.129635.

25. Chen, J., Mohammad, A. & Schedl, T. Comparison of the efficiency of TIR1 transgenes to provoke auxin induced LAG-1 degradation in Caenorhabditis elegans germline stem cells. MicroPubl Biol 2020, (2020).

26. Albarqi, M. M. Y. & Ryder, S. P. The role of RNA-binding proteins in orchestrating germline development in Caenorhabditis elegans. Front. Cell Dev. Biol. 10, 1094295 (2023).

27. Jordan, J. M. et al. Insulin/IGF Signaling and Vitellogenin Provisioning Mediate Intergenerational Adaptation to Nutrient Stress. Current Biology 29, 2380–2388.e5 (2019).

28. Venz, R., Pekec, T., Katic, I., Ciosk, R. & Ewald, C. Y. End-of-life targeted degradation of DAF-2 insulin/IGF-1 receptor promotes longevity free from growth-related pathologies. eLife 10, e71335 (2021).

29. González, A. & Hall, M. N. Nutrient sensing and TOR signaling in yeast and mammals. EMBO J 36, 397–408 (2017).

30. Liu, G. Y. & Sabatini, D. M. mTOR at the nexus of nutrition, growth, ageing and disease. Nat Rev Mol Cell Biol 21, 183–203 (2020).

31. Blackwell, T. K., Sewell, A. K., Wu, Z. & Han, M. TOR Signaling in Caenorhabditis elegans Development, Metabolism, and Aging. Genetics 213, 329–360 (2019).

32. Schreiber, M. A., Pierce-Shimomura, J. T., Chan, S., Parry, D. & McIntire, S. L. Manipulation of behavioral decline in Caenorhabditis elegans with the Rag GTPase raga-1. PLoS Genet 6, e1000972 (2010).

33. Sancak, Y. et al. The Rag GTPases bind raptor and mediate amino acid signaling to mTORC1. Science 320, 1496–1501 (2008).

34. Stojanovski, K. et al. Maintenance of appropriate size scaling of the C. elegans pharynx by YAP-1. Nat Commun 14, 7564 (2023).

35. Sommer, R. J. Phenotypic Plasticity: From Theory and Genetics to Current and Future Challenges. Genetics 215, 1–13 (2020).

36. Minkina, O. & Hunter, C. P. Intergenerational Transmission of Gene Regulatory Information in Caenorhabditis elegans. Trends in Genetics 34, 54–64 (2018).

37. Cenik, E. S. et al. Maternal Ribosomes Are Sufficient for Tissue Diversification during Embryonic Development in C. elegans. Developmental Cell 48, 811–826.e6 (2019).

38. Hibshman, J. D., Webster, A. K. & Baugh, L. R. Liquid-culture protocols for synchronous starvation, growth, dauer formation, and dietary restriction of Caenorhabditis elegans. STAR Protocols 2, 100276 (2021).

39. Ritchie, M. E. et al. limma powers differential expression analyses for RNA-sequencing and microarray studies. Nucleic Acids Res 43, e47 (2015).

40. Huber, W., von Heydebreck, A., Sültmann, H., Poustka, A. & Vingron, M. Variance stabilization applied to microarray data calibration and to the quantification of differential expression. Bioinformatics 18 Suppl 1, S96–104 (2002).

41. Perez-Riverol, Y. et al. The PRIDE database at 20 years: 2025 update. Nucleic Acids Res 53, D543–D553 (2025).

42. Turek, M., Besseling, J. & Bringmann, H. Agarose Microchambers for Long-term Calcium Imaging of Caenorhabditis elegans. JoVE 52742 (2015) doi:10.3791/52742-v.

