## Supplemental Information for "Intergenerational control of ribosomes under dietary restriction"

**Content:**

- Supplemental Figures 1-4
- Supplemental Tables 1-6

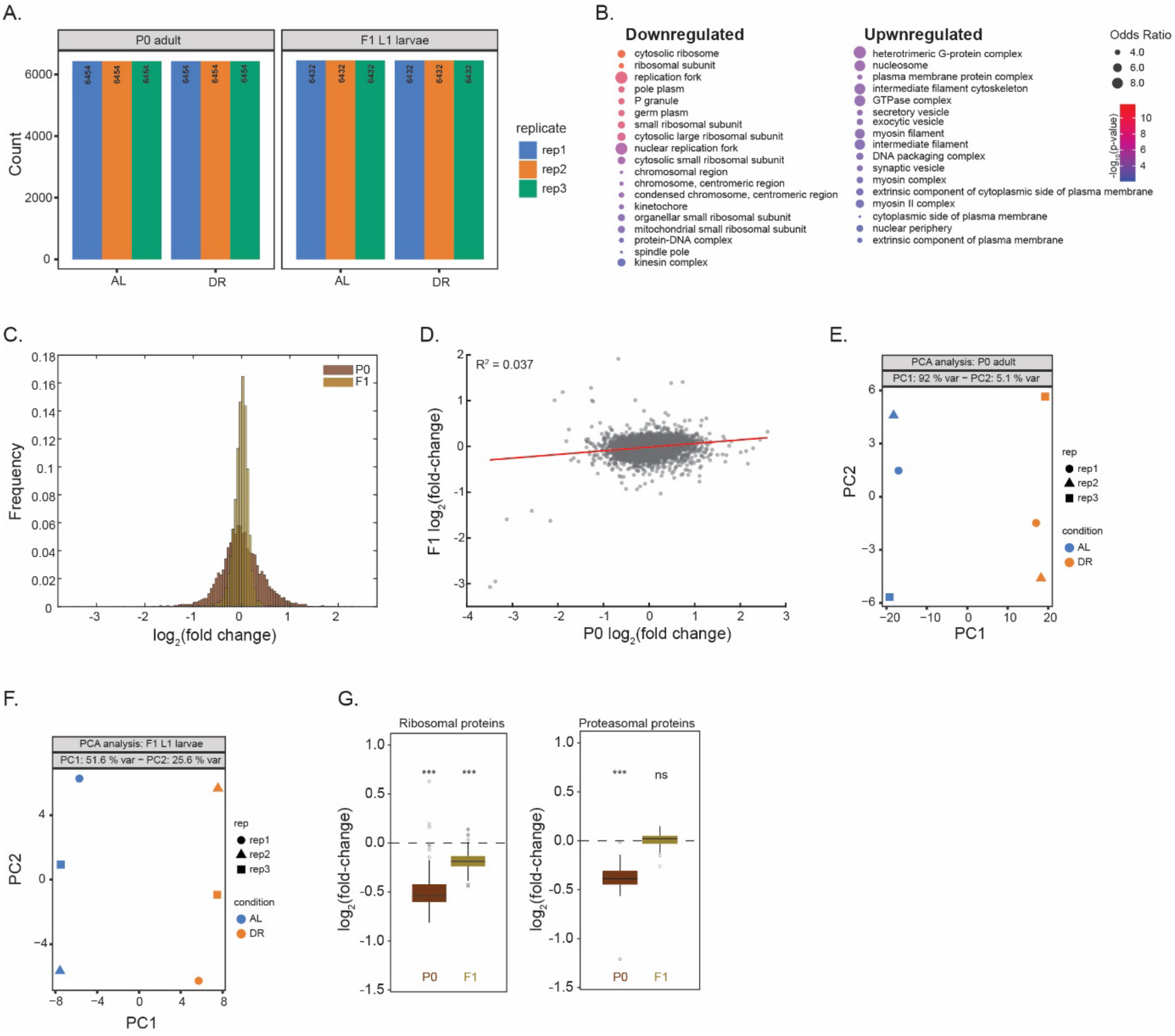


**Supplemental Figure S1. Global proteome changes in response to dietary restriction within and across generations**

**(A)** Number of proteins detected across biological replicates in P0 adults (6454) and F1 L1 larvae (6432) under ad libitum (AL) and dietary restriction (DR) conditions for 3 technical repeats. **(B)** Gene Ontology cellular component terms enriched among significantly downregulated (left) and upregulated (right) proteins in P0 adults under DR compared to AL conditions (FDR ≤ 0.05, minimum 2-fold change). Circle size indicates odds ratio and colour intensity indicates significance, using an enrichment cutoff of odds ratio > 3 and adjusted p-value < 0.01 (Benjamini-Hochberg correction). **(C)** Histograms comparing $log_2$(fold change) in protein abundance between DR and AL conditions for P0 adults (blue) and F1 L1 larvae (orange). P0 adults show a broader distribution of fold changes (wider histogram) compared to F1 larvae, demonstrating larger proteome changes in the parental generation than in their progeny (mean absolute fold change = 0.3 in P0 vs 0.11 in F1; variance = 0.078 in P0 vs 0.016 in F1). **(D)** Correlation between protein fold changes in P0 adults versus F1 L1 larvae ($R^2$ = 0.037), demonstrating weak inheritance of proteome changes across generations. **(E)** Principal component analysis (PCA) of protein abundance data from P0 adults, showing separation between AL and DR conditions across 3 replicates. **(F)** PCA of protein abundance data from F1 L1 larvae showing separation between progeny of AL and DR parents across 3 replicates. **(G)** Box plots showing $log_2$(fold change) in abundance of ribosomal proteins (left) and proteasomal proteins (right) under DR conditions in P0 adults and F1 L1 larvae to respective AL conditions. *** indicates $p < 0.001$.

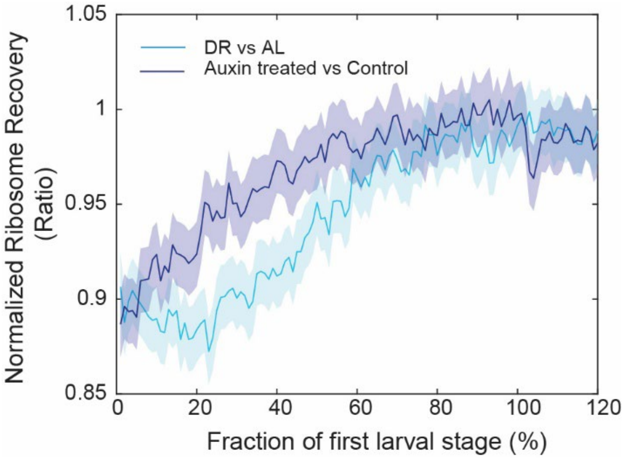


**Supplemental Figure S2. Ribosome recovery in auxin-induced ribosome depleted progeny and DR progeny**

Ribosome levels relative to control during L1 development after ribosome depletion in the maternal proximal germline (dark blue), or after maternal DR (light blue). A value of 1 indicate full recovery of ribosome levels compared to control animals. The recovery occurs slightly faster after auxin-induced ribosome depletion than after maternal DR. Solid lines: mean, shaded regions: 95% confidence interval.

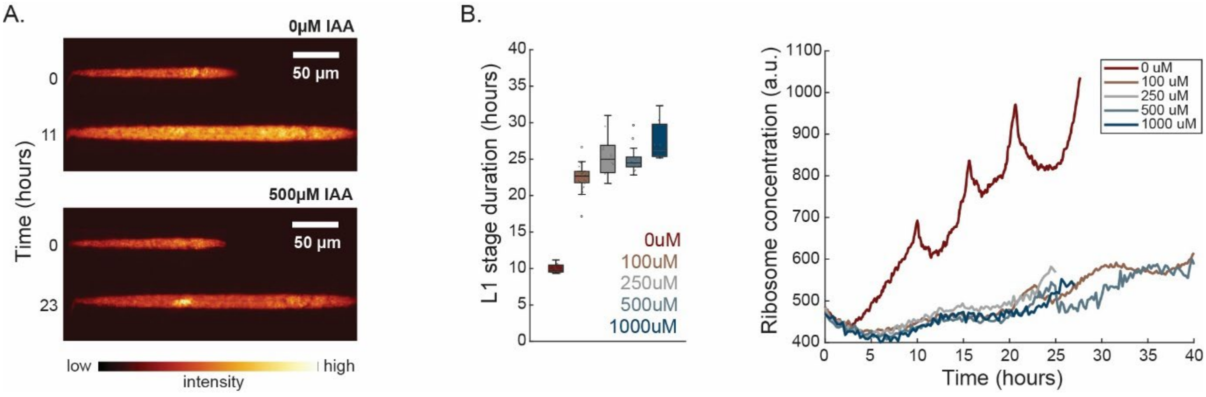


**Supplemental Figure S3. Somatic ribosome depletion causes developmental arrest.**

**(A)** Fluorescent microscopy images of *rpl-34:mcherry, rpl-22-aid, eft-3p:tir-1* animals growing in agarose chambers with 0µM or 500µM IAA immediately after hatching (top) or at indicated time after hatching and IAA exposure (bottom). IAA-treated animals grew more slowly and eventually arrested in development. Images show size matched animals of treated and control group. Bright region in IAA-treated animal corresponds to the germline. Scale bar = 100 µm. Colour scale indicates fluorescence intensity from low (black) to high (yellow). **(B)** Left: L1 stage duration under different auxin concentrations (n≥20 animals per condition). central line: median, box: interquartile ranges (IQR), whisker: ranges except extreme outliers (>1.5*IQR), individual values: crosses, extreme outliers: circles. Development did not proceed to L2 after ribosome depletion. Right: Quantification of RPL-34:mCherry (fluorescence per pixel) over time at indicated auxin concentrations.

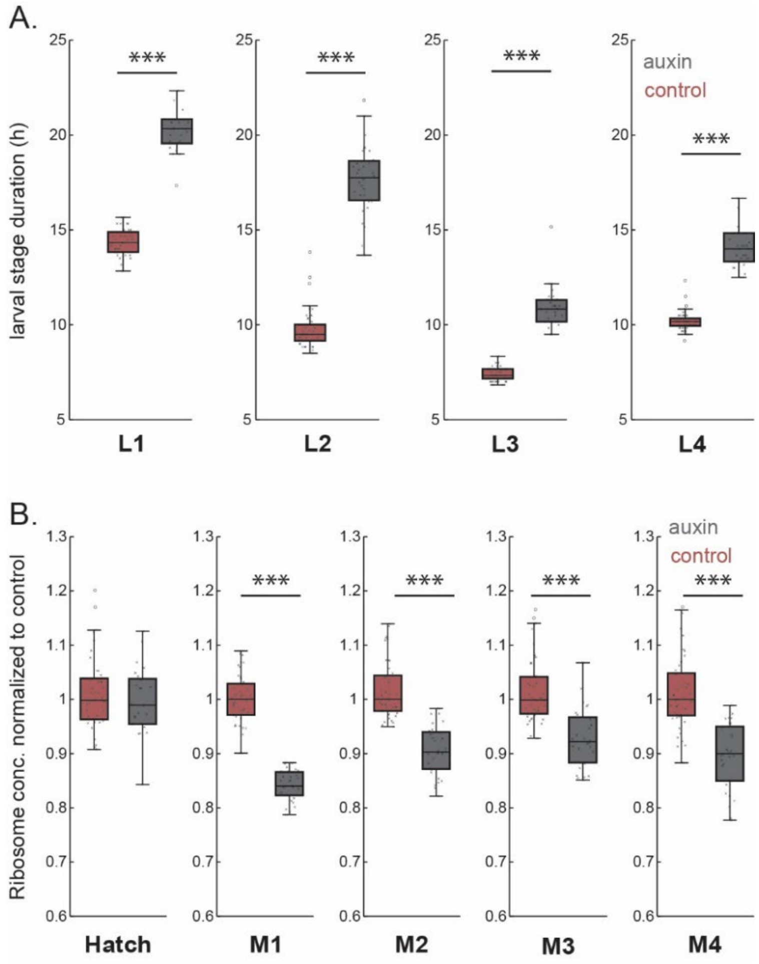


**Supplemental Figure S4. Somatic depletion of RAGA-1 by AID reduces ribosome expression and delays growth**

**(A)** Larval stage duration of *raga-1:gfp:aid; eft-3p:tir-1; rpl-34:mCherry* strain treated with 500 µM auxin (grey) and control (red). central line: median, box: interquartile ranges (IQR), whisker: ranges except extreme outliers (>1.5*IQR), individual values: crosses, extreme outliers: circles. For each condition a total of at least n = 150 individuals were measured on at least m = 3 days. See Supplemental Table S6 for precise sample size and p-values. *** indicate $p<10^{-5}$ (Wilcoxon rank sum test). **(B)** As (A), but for RPL-34:mCherry concentration (intensity per pixel normalized to control).

**Supplemental Table S1a**

Significantly differentially regulated individual proteins after dietary restriction in adults with |$Log_2$(FC)| > 1 and at FDR < 0.05. See separate supplemental file: P0_hits.csv

**Supplemental Table S1b.**

Significantly differentially regulated individual proteins after maternal DR with |$Log_2$(FC)| > 1 and at FDR < 0.05

| Gene | Protein ID | Protein Description | $Log_2$(FC) | FDR |
|---|---|---|---|---|
| CELE_T22B7.3 | Q23042 | Amidinotransferase | 1.407935 | 0.020056 |
| txdc-12.2 | Q9N5S7 | Thioredoxin domain-containing protein | -1.62846 | 0.02811 |
| vit-4 | P18947 | Vitellogenin-4 | -3.07479 | 0.038069 |

**Supplemental Table S2.**

Number of individuals and p-value of comparisons for data shown in Figure 1F

| Measurement | Condition#1 | Condition#2 | n #1 | n #2 | p-value |
|---|---|---|---|---|---|
| RPL-29:GFP | AL P0 | DR P0 | 157 | 160 | $9.4673*10^{-32}$ |
| RPL-29:GFP | AL F1 | DR F1 | 295 | 232 | $3.3715*10^{-34}$ |
| RPL-34:mCherry | AL P0 | DR P0 | 110 | 123 | $1.4356*10^{-27}$ |
| RPL-34:mCherry | AL F1 | DR F1 | 255 | 282 | $8.6893*10^{-17}$ |
| RPN-9:mKate | AL P0 | DR P0 | 248 | 228 | $3.4635*10^{-26}$ |
| RPN-9:mKate | AL F1 | DR F1 | 325 | 329 | $5.5856*10^{-20}$ |

**Supplemental Table S3**

Number of individuals and p-value of comparisons for data shown in Figure 2B-D

| Measurement | Condition#1 | Condition#2 | n #1 | n #2 | p-value |
|---|---|---|---|---|---|
| Ribosomes –hatch | AL F1 | DR F1 | 169 | 170 | $1.01*10^{-33}$ |
| Ribosomes – M1 | AL F1 | DR F1 | 187 | 182 | 0.00721 |
| Ribosomes – M2 | AL F1 | DR F1 | 186 | 181 | 0.0223 |
| Ribosomes – M3 | AL F1 | DR F1 | 174 | 179 | 0.16 |
| Ribosomes – M4 | AL F1 | DR F1 | 174 | 179 | 0.0468 |
| Volume – hatch | AL F1 | DR F1 | 176 | 171 | $6.81*10^{-37}$ |
| Volume – M1 | AL F1 | DR F1 | 188 | 182 | 0.000451 |
| Volume – M2 | AL F1 | DR F1 | 187 | 182 | 0.195 |
| Volume – M3 | AL F1 | DR F1 | 187 | 181 | 0.978 |
| Volume – M4 | AL F1 | DR F1 | 175 | 179 | 0.785 |
| L1 larval duration | AL F1 | DR F1 | 171 | 170 | $1.68*10^{-44}$ |
| L2 larval duration | AL F1 | DR F1 | 187 | 182 | $1.32*10^{-5}$ |
| L3 larval duration | AL F1 | DR F1 | 186 | 181 | 0.0025 |
| L4 larval duration | AL F1 | DR F1 | 174 | 179 | 0.279 |

**Supplemental Table S4**

Number of individuals and p-value of comparisons for data shown in Figure 5E-H

| Measurement | Condition#1 | Condition#2 | n #1 | n #2 | p-value |
|---|---|---|---|---|---|
| Ribosomes – hatch | WT AL - F1 | WT DR - F1 | 118 | 111 | $3.21*10^{-20}$ |
| Ribosomes – hatch | *daf-16* AL - F1 | *daf-16* DR - F1 | 173 | 100 | $5.23*10^{-18}$ |
| L1 duration | WT AL - F1 | WT DR - F1 | 121 | 111 | $4.48*10^{-27}$ |
| L1 duration | *daf-16* AL - F1 | *daf-16* DR - F1 | 173 | 102 | $5.12*10^{-9}$ |
| Volume – hatch | WT AL - F1 | WT DR - F1 | 122 | 116 | $2.69*10^{-25}$ |
| Volume – hatch | *daf-16* AL - F1 | *daf-16* DR - F1 | 174 | 112 | $1.52*10^{-8}$ |

**Supplemental Table S5**

Number of individuals and p-value of comparisons for data shown in Figure 6

| Measurement | Condition#1 | Condition#2 | n #1 | n #2 | p-value |
|---|---|---|---|---|---|
| Ribosomes – hatch | Control - F1 | Parental auxin - F1 | 150 | 218 | $2.03*10^{-32}$ |
| Ribosomes – M1 | Control - F1 | Parental auxin - F1 | 186 | 223 | 0.0005 |
| Ribosomes – M2 | Control - F1 | Parental auxin - F1 | 185 | 221 | 0.000509 |
| Ribosomes – M3 | Control - F1 | Parental auxin - F1 | 183 | 217 | 0.883 |
| Ribosomes – M4 | Control - F1 | Parental auxin - F1 | 183 | 217 | 0.127 |
| Volume – hatch | Control - F1 | Parental auxin - F1 | 151 | 218 | $6.81*10^{-37}$ |
| Volume – M1 | Control - F1 | Parental auxin - F1 | 186 | 225 | 0.000451 |
| Volume – M2 | Control - F1 | Parental auxin - F1 | 187 | 223 | 0.195 |
| Volume – M3 | Control - F1 | Parental auxin - F1 | 185 | 222 | 0.978 |
| Volume – M4 | Control - F1 | Parental auxin - F1 | 183 | 218 | 0.785 |
| L1 larval duration | Control - F1 | Parental auxin - F1 | 151 | 218 | $1.06*10^{-48}$ |
| L2 larval duration | Control - F1 | Parental auxin - F1 | 186 | 223 | 0.88 |
| L3 larval duration | Control - F1 | Parental auxin - F1 | 185 | 221 | $4.14*10^{-13}$ |
| L4 larval duration | Control - F1 | Parental auxin - F1 | 183 | 217 | 0.00088 |

**Supplemental Table S6**

Number of individuals and p-value of comparisons for data shown in Figure S4

| Measurement | Condition#1 | Condition#2 | n #1 | n #2 | p-value |
|---|---|---|---|---|---|
| L1 larval duration | Control - P0 | Auxin tr. - P0 | 41 | 25 | $1.06*10^{-10}$ |
| L2 larval duration | Control - P0 | Auxin tr. - P0 | 46 | 31 | $1.38*10^{-13}$ |
| L3 larval duration | Control - P0 | Auxin tr. - P0 | 46 | 31 | $1.12*10^{-13}$ |
| L4 larval duration | Control - P0 | Auxin tr. - P0 | 45 | 30 | $2.6*10^{-13}$ |
| Ribosomes - hatch | Control - P0 | Auxin tr. - P0 | 45 | 30 | 0.597 |
| Ribosomes – M1 | Control - P0 | Auxin tr. - P0 | 41 | 25 | $1.36*10^{-13}$ |
| Ribosomes – M2 | Control - P0 | Auxin tr. - P0 | 46 | 31 | $2.28*10^{-12}$ |
| Ribosomes – M3 | Control - P0 | Auxin tr. - P0 | 46 | 31 | $3.59*10^{-8}$ |
| Ribosomes – M4 | Control - P0 | Auxin tr. - P0 | 45 | 30 | $6.84*10^{-10}$ |
